# Genetic, structural, and functional analysis of mutations causing methylmalonyl-CoA epimerase deficiency

**DOI:** 10.1101/484048

**Authors:** Kathrin Heuberger, Henry J. Bailey, Patricie Burda, Apirat Chaikuad, Ewelina Krysztofinska, Terttu Suormala, Céline Bürer, Seraina Lutz, Brain Fowler, D. Sean Froese, Wyatt W. Yue, Matthias R. Baumgartner

**Author notes:** equal contribution (Kathrin Heuberger, Henry J. Bailey). Corresponding authors: Wyatt W. Yue and Matthias R. Baumgartner. Author contributions - D.S.F., B.F., M.R.B. and W.W.Y conceived of the project. D.S.F., P.B. and W.W.Y. designed the research. P.B., D.S.F. and W.W.Y. analysed the results. D.S.F. performed the size exclusion chromatography. K.H. performed the Western blots and [^14^C]-propionyl-CoA enzymatic assay. H.J.B. performed biophysical characterization of MCEE variants. C.B. sequenced the patients and identified the mutations. S.L. and T.S. performed the propionic acid incorporation assay. D.S.F., E.K. and H.J.B. produced the recombinant proteins. A.C. and H.J.B. solved the crystal structures. D.S.F. and W.W.Y. wrote the paper with contributions from other authors.

## Abstract

Human methylmalonyl-CoA epimerase (*MCEE*) catalyzes the interconversion of D-methylmalonyl-CoA and L-methylmalonyl-CoA in propionate catabolism. Autosomal recessive mutations in *MCEE* reportedly cause methylmalonic aciduria (MMAuria) in eleven patients. We investigated a cohort of 150 individuals suffering from MMAuria of unknown origin, identifying ten new patients with mutations in *MCEE*. Nine patients were homozygous for the known nonsense mutation p.Arg47* (c.139C>T), and one for the novel missense mutation p.Ile53Arg (c.158T>G). To understand better the molecular basis of MCEE deficiency, we mapped p.Ile53Arg, and two previously described patient mutations p.Lys60Gln and p.Arg143Cys, onto our 1.8 Å structure of wild-type (wt) human MCEE. This revealed potential dimeric assembly disruption by p.Ile53Arg, but no clear defects from p.Lys60Gln or p.Arg143Cys. Functional analysis of MCEE-Ile53Arg expressed in a bacterial recombinant system as well as patient-derived fibroblasts revealed nearly undetectable soluble protein levels, defective globular protein behavior, and using a newly developed assay, lack of enzymatic activity - consistent with misfolded protein. By contrast, soluble protein levels, unfolding characteristics and activity of MCEE-Lys60Gln were comparable to wt, leaving unclear how this mutation may cause disease. MCEE-Arg143Cys was detectable at comparable levels to wt MCEE, but had slightly altered unfolding kinetics and greatly reduced activity. We solved the structure of MCEE-Arg143Cys to 1.9 Å and found significant disruption of two important loop structures, potentially impacting surface features as well as the active-site pocket. These studies reveal ten new patients with MCEE deficiency and rationalize misfolding and loss of activity as molecular defects in MCEE-type MMAuria.

## Introduction

Propionyl-CoA is the common degradation product from branched-chain amino acids, odd-chain fatty acids, and the side chain of cholesterol. The propionate catabolic pathway serves to funnel propionyl-CoA into the tricarboxylic acid (TCA) cycle for use as cellular energy sources through oxidative phosphorylation. Located at the centre of this pathway, methylmalonyl CoA epimerase (MCEE) catalyzes the epimerization of D-methylmalonyl-CoA, generated from propionyl-CoA by propionyl-CoA carboxylase (PCC), to form L-methylmalonyl-CoA, subsequently converted into succinyl-CoA by methylmalonyl-CoA mutase (MUT) for entry into the TCA cycle.

Isolated methylmalonic aciduria (MMAuria), an inborn error of organic acid metabolism, is typically caused by deficiency of MUT or by a defect in the transport or processing of its organometallic cofactor, adenosylcobalamin. However, mutations in the human *MCEE* gene (OMIM #251120) have been identified in eleven cases of atypical MMAuria (1-6). For two patients, coincidental mutations in the *SPR* gene causing sepiapterin reductase deficiency sufficiently explained their clinical symptoms (2,4), while two others have been described as asymptomatic (3), leaving the clinical importance of MCEE deficiency in doubt. The majority of patients with MCEE deficiency (seven including those with sepiapterin reductase deficiency) are homozygous for the stop-gain nonsense mutation c.139C>T (p.Arg47*)(1-6); the missense mutations p.Lys60Gln and p.Arg143Cys (3) and a splicing mutation (c.379-644A>G) (5) have also been identified, but the functional relevance of these missense mutations remains unclear.

In the human genome, MCEE is one of six proteins belonging to the vicinal oxygen chelate (VOC) superfamily, which include also glyoxalase I (*GLO1* gene, GLOD1 protein), 4-hydroxyphenylpyruvic acid dioxygenase (*HPD*, GLOD3), 4-hydroxyphenylpyruvic acid dioxygenase-like (*HPDL*, GLOXD1), and glyoxalase domain-containing 4 (*GLOD4*) and 5 (*GLOD5*). VOC members are metalloenzymes highly divergent in sequence and biological functions, but universally share the use of the βαβββ structural motif (also known as the glyoxylase fold) to build a divalent metal-containing active site (7,8). VOC enzymes catalyze a range of chemical reactions including isomerization, epimerization, oxidative C-C bond cleavage and nucleophilic substitution (9). The active-site divalent metal is used to bind the reaction substrate, intermediates or transition states in a bidentate fashion (8). To date, only structures from bacterial MCEE orthologues (*Propionibacterium shermanii* and *Thermoanaerobacter tengcongensis*) have been reported (10,11).

In this article, we conducted an in depth investigation of MCEE deficiency at the gene and protein levels. From a cohort of 150 patients with MMAuria of unknown etiology, we identified ten new patients with mutations on the *MCEE* gene including a novel missense mutation. We determined the crystal structure of human MCEE of the wild-type as well as p.Arg143Cys variant proteins. We further characterized protein expression and enzyme activity for the three known MCEE missense mutations associated with disease. Our study provides a molecular explanation for the biochemical defects associated with the missense mutations.

## Results & Discussion

### Identification of ten new MCEE patients with methylmalonic aciduria

Thus far, 11 cases of MMAuria have been identified due to mutations in the *MCEE* gene (1-6). We screened fibroblast cell lines taken from 150 patients with mild but clear MMAuria who could not be assigned to a cobalamin class of defect. From this cohort, we identified ten patients from nine families with mutations in *MCEE* (Table 1). All patients are homozygous for their respective mutations, of which nine out of ten harbour the previously described c.139C>T (p.Arg47*) nonsense mutation. This remains by far the most common mutation identified in MCEE deficiency, with 16 out of 21 patients homozygous for this allele. One patient in our cohort harboured c.158T>G (p.Ile53Arg), a novel missense mutation that is not found in the ExAC database (>120,000 alleles) (12). We did not observe either c.178A<C (p.Lys60Gln), previously found in one patient in the homozygous state, or c.427C>T (p.Arg143Cys), previously found in two patients in the heterozygous state without an apparent second mutation (3).

**Table 1.** List of ten newly identified MCEE patients with relevant genetic, biochemical and clinical data.

| Patient No. | Age at onset | Mutation<sup>a</sup> | Clinical presentation, laboratory data | Urinary MMA<br>mmol/mol creat.<br>Ref. 0.3 – 1.1 | Propionate incorporation<sup>b</sup><br>nmol/16h/mg |  |
| --- | --- | --- | --- | --- | --- | --- |
|  |  |  |  |  | - OHCbl<br>Ref. 3.5 – 24.4 | + OHCbl<br>Ref. 4.3 – 28.9 |
|  |  | homozygous for all patients |  |  |  |  |
| 1 | 4 mo | c.139C>T; p. Arg47* | Cardiomyopathy. Sibling with similar biochemical profile but no symptoms | 441 | 1.9 (1.8 – 2.0) | 1.9 (1.8 – 2.1) |
| 2 | 2 yr | c.139C>T; p. Arg47* | Severe metabolic acidosis and hypoglycemia following gastroenteritis, elevated propionyl-carnitine; 3-OH propionate & methyl-citrate in urine | 458 | 1.9 (1.8 - 2.1) | 2.0 (1.8 – 2.1) |
| 3 | 2.5 yr | c.139C>T; p. Arg47* | Severe metabolic acidosis, elevated propionyl-carnitine; 3-OH propionate & methyl-citrate | elevated | 1.4 (1.2 - 1.5) | 1.4 (1.2 – 1.6) |
| 4 | 1.5 yr | c.139C>T; p. Arg47* | Severe metabolic acidosis and hypoglycemia following intercurrent illness | 594 | 2.0 (1.8 - 2.3) | 1.9 (1.5 – 2.4) |
| 5<sup>c</sup> | - | c.139C>T; p. Arg47* | No data; enzyme assays performed at 3 years of age | elevated | 3.0 (2.9 - 3.0) | 3.0 (2.9 – 3.4) |
| 6<sup>c</sup> | - | c.139C>T; p. Arg47* | No data; enzyme assays performed at 7 years of age | elevated | 2.7 (2.6 – 3.0) | 2.7 (2.4 – 2.9) |
| 7 | 2 yr | c.139C>T; p. Arg47* | Slow motor development, hypotonia (inability to walk independently), spasticity of legs, eczema, episode of vomiting and diarrhea | 143 | 2.5 (2.2 - 2.8) | 2.5 (2.3 – 2.7) |
| 8 | 1 mo | c.139C>T; p. Arg47* | Sepsis, psychomotor retardation, seizures, elevated C3-acylcarnitine, methyl-citrate in urine | elevated | 2.0 (1.8 - 2.1) | 2.0 (1.9 – 2.2) |
| 9 | 6 mo | c.158T>G; p.Ile53Arg | Seizures and hypoglycemia following viral infection, elevated propionyl-carnitine; methyl-citrate in urine; then normal development | 143-184 | 3.0 (2.9 - 3.2) | 3.0 (2.9 – 3.1) |
| 10 | < 1 yr | c.139C>T; p.Arg47* | Ketoacidosis | elevated | 2.4 (2.4 - 2.3) | 2.4 (2.4 - 2.4) |

<sup>a</sup> According to NM\_032601 and NP\_1159990. Nucleotide numbering uses +1 as the A of the ATG translation initiation codon in the reference sequence, with the initiation codon as codon 1.
<sup>b</sup> [<sup>14</sup>C]propionate incorporation: fibroblasts were grown for 3 days without (-) and with (+) 10mg of hydroxocobalamin (OH-Cbl)/ml medium. Values for patient cells represent mean and range (in brackets) from 3 replicate experiments. For controls (WT) the range of 33 individual fibroblast cell lines is given.
<sup>c</sup> Patient 5 and patient 6 are siblings.

In our cohort, disease onset ranged from 1 month old to 2.5 years of age, while from two patients we had no information. Clinical symptoms were variable but usually mild, and no patient was responsive to vitamin B_12_ treatment. At least three patients presented following an intercurrent illness, while three presented with metabolic acidosis and/or hypoglycemia. In addition, elevated levels of other metabolites typical for a block in the propionate degradation pathway, such as 2-methylcitrate, propionylcarnitine and 3-hydroxypropionate, were documented in most patients. Investigation in patient fibroblasts revealed mildly decreased propionate incorporation (Table 1), which did not increase upon addition of hydroxocobalamin.

### Structural features of human MCEE

We performed structural biology studies to establish the molecular basis of disease-causing mutations on the human MCEE protein (hMCEE). As a first step, we determined the crystal structure of wild-type (wt) protein (hMCEE_WT_) to 1.8 Å resolution (Table 2), as part of a wider effort that also generated crystal structures of three other human VOCs, namely hHPD (PDB: 3ISQ), hGLOD4 (PDB 3ZI1) and hGLOD5 (PDB 3ZW5). Most VOC members are structurally made up of four glyoxylase (GLOD) motifs of βαβββ topology. As exemplified in Supplemental Fig. S1, the human VOCs display versatility in the way that four GLOD motifs are assembled, at the gene or protein level, to give rise to a minimal functional unit harbouring two metal-binding active sites.

**Table 2.** Data collection and refinement statistics for the crystal structures. Data from highest resolution shell shown in parenthesis.

| Dataset | hMCEE<sub>WT</sub> | hMCEE<sub>R143C</sub> |
| --- | --- | --- |
| Beamline | Diamond Beamline I02 | Diamond Beamline I04 |
| Wavelength (Å) | 0.9763 | 0.9795 |
| Unit cell parameters (a,b,c) | 53.12 Å, 66.69 Å, 76.98 Å | 53.16 Å, 66.99 Å, 77.13 Å |
| ($\alpha$, $\beta$, $\gamma$) | 90.0°, 90.1°, 90.0° | 90.00°, 90.02°, 90.00° |
| Space group | P 1 21 1 | P 1 21 1 |
| Resolution range (Å) | 36.58 – 1.80 | 47.77 – 1.93<sup>*</sup> |
| Observed/Unique reflections | 49519 | 19630 |
| R<sub>sym</sub> (%) | 0.076 (0.56) | 0.13 (0.92) |
| I/$\sigma$(I) | 1.98 | 7.2 (1.2) |
| Completeness (%) | 99.2 (99.7) | 88.9 (55.8) |
| Multiplicity | 3.6 (3.6) | 3.3 (2.7) |
| R<sub>cryst</sub> (%) | 0.19 | 0.23 |
| R<sub>free</sub> (%) | 0.238 | 0.28 |
| Wilson B factor (Å<sup>2</sup>) | 23.7 | 25 |
| Average total B factor (Å<sup>2</sup>) | 27.59 | 29.46 |
| R.m.s.d. bond length (Å) | 0.67 | 0.008 |
| R.m.s.d. bond angle (°) | 0.69 | 1.070 |
| Missing residues | 1-44 | 1-44 |
| Clashscore | 6 | 5.78 |
| Ramachadran favoured (%) | 97 | 97 |
| Ramachandran disallowed (%) | 1 | 0.2 |
| Rotamer outliers (%) | 0 | 0.5 |

\* Anisotropic data truncated in staraniso using local I/ $\sigma$ (I) cut off at 1.2 results in the inclusion of data to 1.9 Å with outer-shell ellipsoidal completeness at 58.8% and spherical completeness at 10%.

In the case of hMCEE, two βαβββ motifs pack side-by-side to form an 8-stranded sheet that completes the active site for one protomer (Fig. 1A). Crystallographic (supplemental Fig. S1) and solution data (supplemental Fig. S2) show that hMCEE is dimeric, in agreement with bacterial orthologs (10,11). The VOC members are so called because a divalent metal per active site can bind the substrate, intermediates or transition states in a bidentate fashion (8). The divalent metal ion in hMCEE, observed as cobalt in our structure, is coordinated by three residues strictly conserved among VOCs: His50 (strand β1), His122 (strand β5), and Glu172 (strand β8) (Fig. 1B).

**Figure 1.**
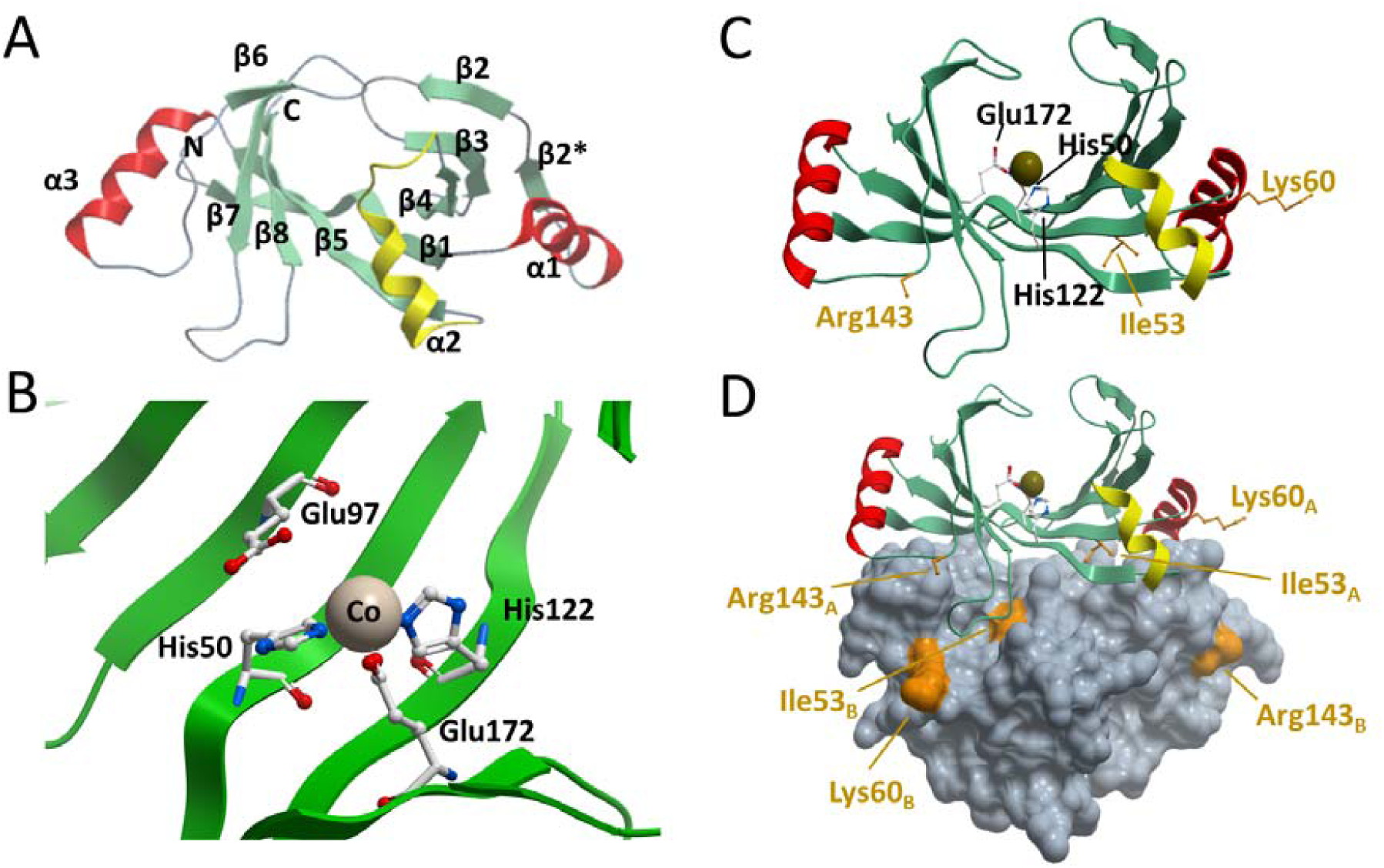
Structure of hMCEE (PDB ID:3RMU). **A.** Monomeric structure coloured by secondary structure, whereby β-strands are green, α-helices that are part of the GLOD motif are red, and regions that connect GLOD motifs are yellow. **B.** Active-site architecture showing the divalent metal and metal coordinating residues. Pink sphere: cobalt. **C.** On the hMCEE monomer, residues that are mutated in disease are labeled in orange, active-site residues are labeled in black. **D.** hMCEE physiological dimer with the second subunit represented as grey space fill. Mutations are labeled in orange and designated as belonging to chain (monomer) A or B.

### Structural mapping of MCEE mutations

Both the reported p.Arg143Cys and newly identified p.Ile53Arg missense mutations are predicted by SIFT (Damaging, score: 0.00 & 0.01) and PolyPhen2 (Probably damaging, score: 0.989 & 1.00) to be deleterious. By contrast, the other previously reported mutation p.Lys60Gln was predicted to be not deleterious (SIFT: 0.12 tolerated, PolyPhen2: 0.04 benign). The most common mutation of MCEE, c.139C>T, causes truncation of the protein at p.Arg47*, within the first β-sheet (supplemental Fig. S3). Loss of almost the entire protein is therefore the likely cause of enzymatic dysfunction due to this mutation, assuming there is residual mRNA following nonsense-mediated decay.

By contrast, the molecular dysfunction due to the missense mutations is less clear (Fig. 1C). In our hMCEE structure, Ile53 is located at the dimeric interface, making hydrophobic contacts with Gly168 and Val169 from the loop region connecting strands β7 and β8 (loop_β7-_ β8) of the other subunit in the dimer, (Fig. 1D). The amino acid (aa) position of Ile53 is highly conserved among MCEE orthologues (85% occupied by Ile, n=150). An Ile-to-Arg at this position likely interferes with proper dimeric assembly, and is predicted by FoldX (13) to have severely reduced stability (ΔΔG 9.53 kcal/mol). By contrast, Lys60 and Arg143 occupy amino acid positions that are more variable. Position 60 is only occupied by lysine in 15% of MCEE homologs while position 143 is 60% occupied by arginine. Both residues are surface exposed and not directly involved in the dimeric interface and active-sites of both subunits (Fig. 1D), consistent with a FoldX prediction of no effect for p.Lys60Gln (ΔΔG 0.3 kcal/mol), and moderately reduced stability for p.Arg143Cys (ΔΔG 2.26 kcal/mol).

### Characterization of MCEE mutations in recombinant and patient cells

To validate our structural interpretation of mutations, we performed expression studies in *E. coli* and human cells. When expressed in *E. coli* (Fig. 2A), hMCEE wt, p.Lys60Gln and p.Arg143Cys proteins were highly soluble, while the p.Ile53Arg variant showed a massive decrease in protein solubility, consistent with a poorly folded protein. All wt and variant hMCEE molecules eluted at similar volumes by size exclusion chromatography (supplemental Fig. S4). Thermal unfolding (Fig. 2B) of purified hMCEE p.Ile53Arg by nano-differential scanning fluorimetry revealed a melting curve of multiple transitions, indicative of heterogenous protein states. By contrast, purified wt and p.Lys60Gln proteins showed globular protein behaviour reflected by a cooperative sigmoidal melting pattern, indicating a single unfolding/folding transition. p.Arg143Cys also behaved similarly to wt and p.Lys60Gln, however late stage unfolding/folding intermediates deviate from sigmoidal melting. Together our data indicate reduced thermostability for the p.Ile53Arg variant protein and a potential slight alteration in that of p.Arg143Cys.

**Figure 2.**
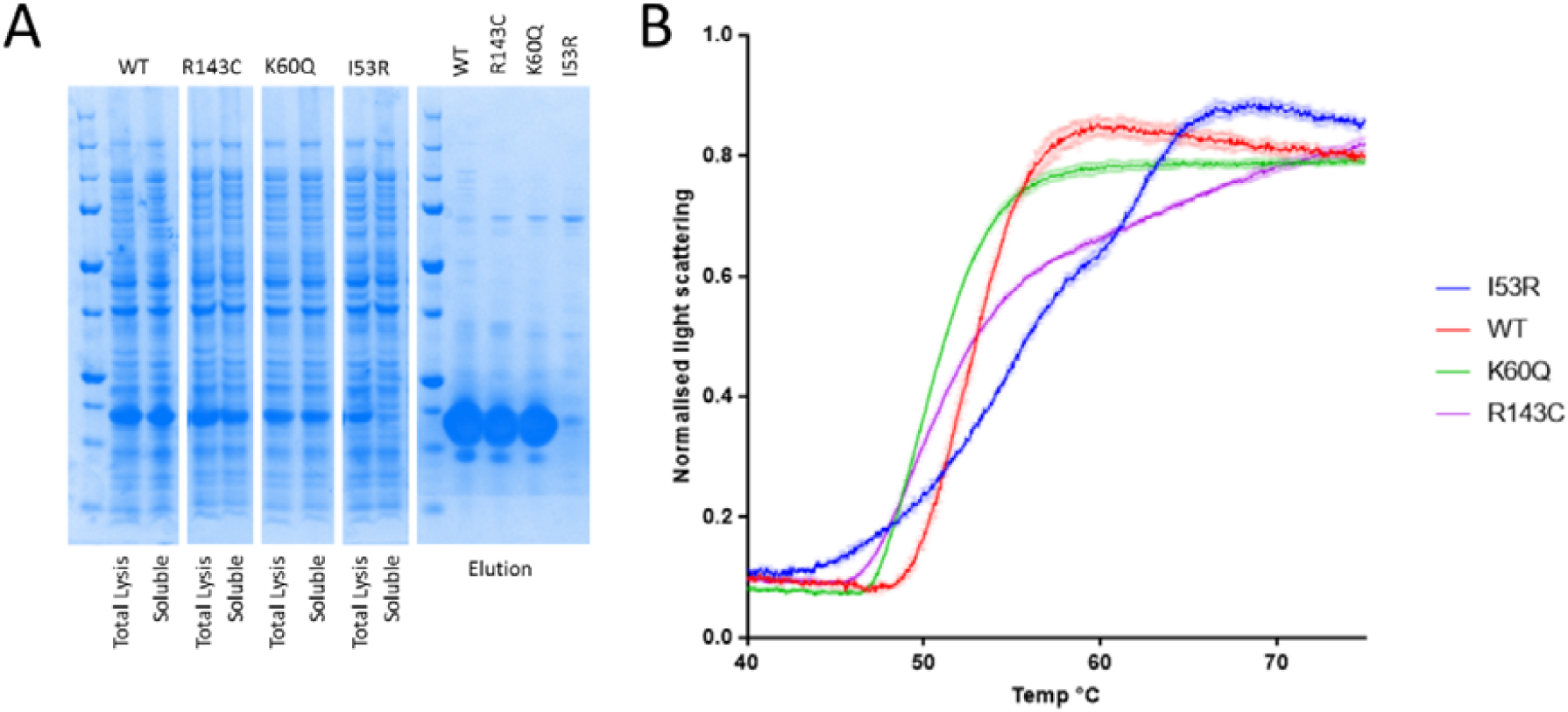
Characterization of recombinant hMCEE wt and variant proteins. **A.** SDS-PAGE analysis of hMCEE protein expression (from cell lysate: Total Lysis; and centrifuged supernatant fractions: Soluble) and solubility (from eluant fractions of affinity purification); **B.** nanoDSF melting curve plot of normalized light scattering intensity against temperature.

These results were validated by over-expression of hMCEE in patient fibroblasts homozygous for the *MCEE* null mutation (c.139C>T; p.Arg47*). For this experiment, flag-tagged wt and mutant hMCEE proteins were over-expressed with visualization of the flag-tag by Western blot analysis (Fig. 3*A*). Expression of each construct performed at least 3 times revealed wt protein to be well expressed, while hMCEE containing p.Lys60Gln and p.Arg143Cys were detectable at only slightly lower levels (63 ± 17% and 74 ± 26% of wt, respectively) (Fig. 3*B*). However, hMCEE containing p.Ile53Arg had very low levels of detectable protein (6 ± 1% of wt), similar to empty vector (4 ± 2% of wt) (Fig. 3*B*). Thus, consistent with recombinant studies, it appears that p.Ile53Arg causes an inability to fold correctly.

**Figure 3.**
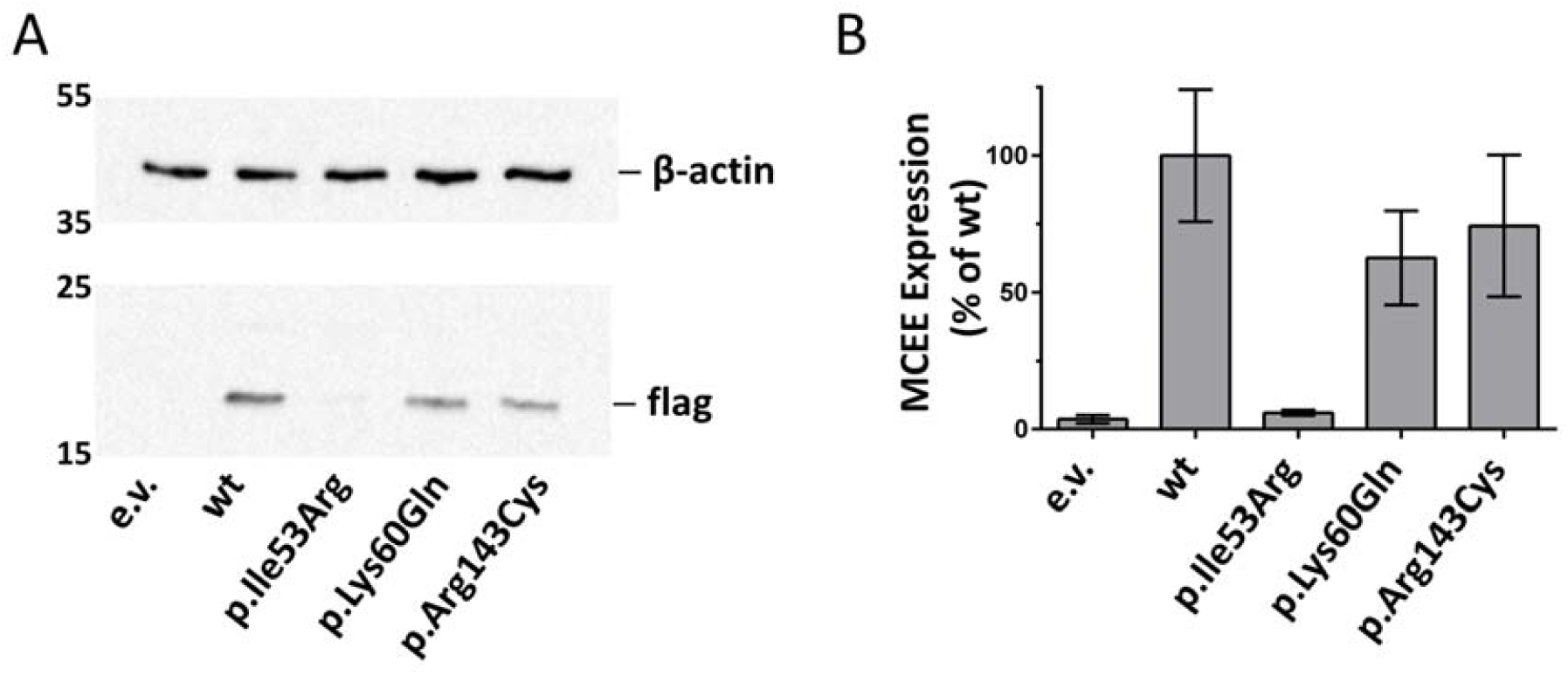
Over-expression of hMCEE-flag in human fibroblasts. **A.** Representative Western blots depicting detection of wild-type (wt) or mutant hMCEE-flag following over-expression in patient fibroblasts deficient for MCEE enzyme. Vector without insert was used as a control (e.v.). Loading was controlled by detection of endogenous β-actin. Numbers on the left correspond to molecular weights (kDa). Approximate expected molecular weights, hMCEE-flag: 18 kDa; β-actin: 42 kDa. **B.** Bar-graph depicting mean and standard deviation of Western blot results performed in 3 independent experiments.

### Structure of human MCEE p.Arg143Cys variant

We determined the crystal structure of the p.Arg143Cys variant protein (hMCEE_R143C_) to 1.9 Å resolution (Table 2), to directly inspect the atomic environment of the substitution (Fig. 4). While hMCEE_R143C_ superimposes well overall with hMCEE_WT_ (Cα-RMSD 0.278 Å), significant main-chain displacement was clearly observed in the loop region connecting helix α3 and β6 (loop_α3-β6_) that harbours the site of mutation at aa 143, as well as the nearby loop_β7-β8_ (Fig. 4, inset). These loop regions connect several β-strands that make up the protomer active site. In the hMCEE_R143C_ structure, the substituted Cys143 residue generated more mobility and disorder within the loop_α3-β6_. As a result, the main-chain atoms of aa 142-146 are displaced by ∼2.5-4.7 Å compared to hMCEE_WT_. The increased mobility in loop_α3-β6_ impacts on the nearby loop_β7-β8_ that packs against it, resulting in 2.0-4.8 Å main-chain displacement at the first half of loop_β7-β8_ (aa 164-167). The second half of loop_β7-β8_, involved in the aforementioned dimeric interface, was not affected by any displacement. Together, these local structural changes have the potential to impact on the surface features of the homodimer, as well as the active site pocket.

**Figure 4.**
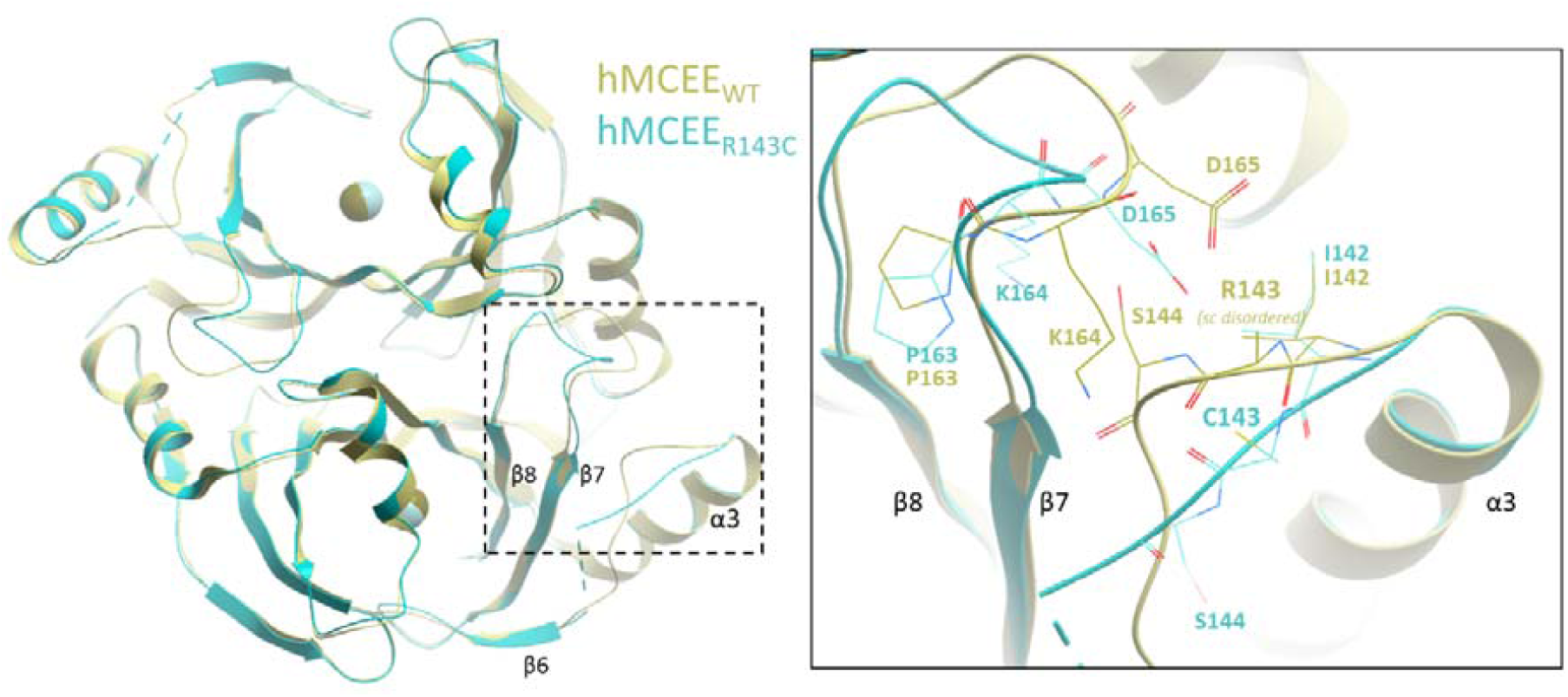
Structure of hMCEE p.Arg143Cys variant. Structural superposition of hMCEE homodimer from hMCEE_WT_ (yellow) and hMCEE_R143_ (cyan) proteins. ***Inset.*** Magnified view of the local atomic environment surrounding the Arg143/Cys143 site, where amino acid side-chains are shown as thin lines.

### Biochemical characterization of MCEE mutations

We developed a radioactive HPLC-based assay to assess MCEE activity of wt and variants. This assay follows the production and separation of [^14^C]-methylmalonic acid and [^14^C]-succinic acid from [^14^C]-propionic acid following addition of [^14^C]-propionyl-CoA to fibroblast cell lysates, as depicted in Fig. 5*A*. Using UV detection, we were able to detect separated propionic acid, methylmalonic acid and succinic acid following HPLC analysis (supplemental Fig. S5). In cell lysates from control fibroblasts, we identified high levels of [^14^C]- methylmalonic acid but only very little [^14^C]-succinic acid (Fig. 5*B*; supplemental Fig. S6*A*). To determine if MUT or MCEE was rate-limiting for succinate production, we expressed each individually, or together (Fig. 5*B*; supplemental Fig. S6*B*-*C*). While over-expressed MUT-alone only marginally increased detectable [^14^C]-succinic acid, MCEE expression alone, and especially co-expression with MUT, provided a marked increase in [^14^C]-succinic acid (Fig. 5*B*; supplemental Fig. 6*D*). Therefore, MCEE appears to be the rate-limiting step in this pathway. This same basic pattern could be seen in MCEE null fibroblasts (Fig. 5*C*).

**Figure 5.**
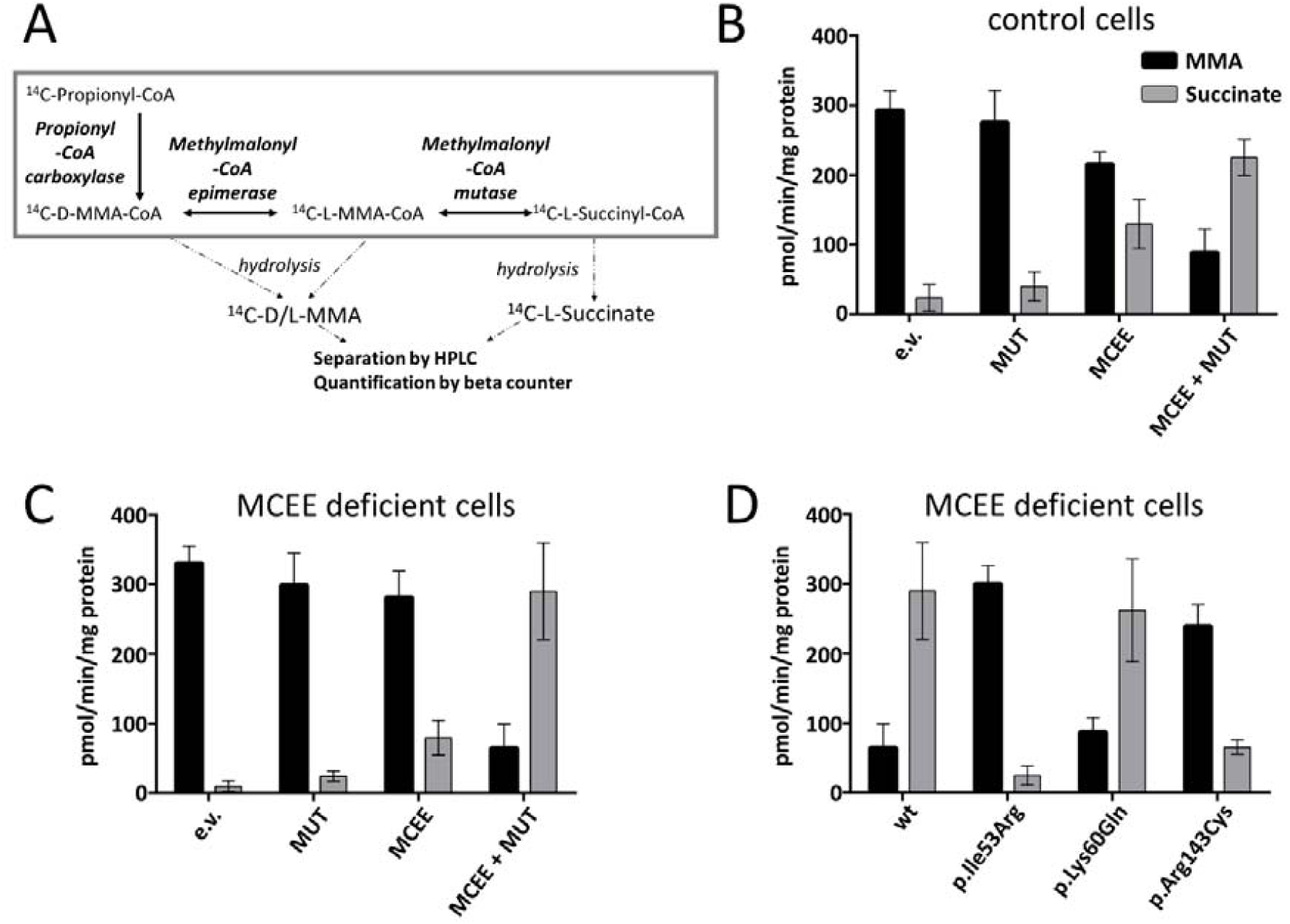
Biochemical analysis of hMCEE disease associated mutations. **A.** Schematic depiction of the cellular pathway leading from propionyl-CoA to succinyl-CoA and the enzymes involved in this process. **B and C.** Bar-graph showing the amount of methylmalonic acid (MMA) and succinate following over-expression of empty vector (e.v.), methylmalonyl-CoA mutase (MUT) alone, MCEE alone, or both MCEE and MUT in control fibroblasts (**B**) and patient fibroblasts deficient in MCEE (**C**). **D.** Bar graph depicting production of MMA and succinate following over-expression of wild-type (wt) and mutant MCEE in the presence of over-expressed MUT in MCEE patient fibroblasts. Results shown represent the mean and standard deviation of 3 independent experiments.

We further examined the effect of the mutant MCEE proteins in the presence of over-expressed MUT in MCEE null fibroblast lysates (Fig. 5*D*). Over-expression of MCEE harbouring p.Ile53Arg resulted in very little detectable [^14^C]-succinate. This is consistent with an inability to convert D-methylmalonyl-CoA to L-methylmalonyl-CoA due to a lack of correctly folded protein, as was demonstrated by Western blot analysis. By contrast, over-expressed MCEE harbouring p.Lys60Gln produced similar levels of detectable succinate as wt protein. Finally, despite being well expressed, MCEE harbouring p.Arg143Cys had markedly reduced [^14^C]-succinic acid production, agreeing with the significantly altered local environment in the hMCEE_R143C_ structure. This conformational change could impair enzymatic activity of p.Arg143Cys either by direct structural interference with the active site, or indirectly e.g. via loss of essential interactions with other proteins in the succinyl-CoA production pathway.

## Conclusions

The identification of an additional ten patients with MCEE deficiency adds more information toward the debate of whether deficiency of this protein does indeed cause disease and not just elevated methylmalonic acid levels, and confirms that complete MCEE deficiency, despite low penetrance, may lead to an acute clinical phenotype with metabolic crisis resembling classical organic acidurias, similar to e.g. 3-methylcrotonyl-CoA carboxylase deficiency (14). Our combined structural, biophysical and enzymatic assessment of MCEE defects confirm the importance of this protein within the propionate degradation pathway. While protein-destabilizing mutations (e.g. p.Ile53Arg) explains enzymatic defects more readily, mutations away from the active-site (e.g. p.Arg143Cys) could still cause a loss of activity, although the underlying mechanism needs further clarification. In regards to p.Lys60Gln, however, we could identify no defect conferred to the protein, leaving the molecular mechanism and its potential to cause disease in doubt.

## Supporting information

## Acknowledgements

This work was supported by the Rare Disease Initiative Zurich, a Clinical Research Priority Program from the University of Zurich (to D. S. F. and M. R. B.) and the Swiss National Science Foundation [SNSF 31003A_175779] (to M.R.B.). The SGC is a registered charity (number 1097737) that receives funds from AbbVie, Bayer Pharma AG, Boehringer Ingelheim, Canada Foundation for Innovation, Eshelman Institute for Innovation, Genome Canada, Innovative Medicines Initiative (EU/EFPIA) [ULTRA-DD grant no. 115766], Janssen, MSD, Merck KGaA, Novartis Pharma AG, Ontario Ministry of Economic Development and Innovation, Pfizer, São Paulo Research Foundation-FAPESP, Takeda and Wellcome Trust [106169/ZZ14/Z]. We thank Dr. Holger Blessing (Universitätsklinikum Erlangen, Germany), Dr. Ernst Christensen (Rigshospitalet, Copenhagen, Denmark), Dr. Andrew Morris (Alder Hey, Liverpool, UK), Dr. Murray Bain (St. George’s Hospital Medical School, London, UK), Dr. Mike Champion (St. Thomas’ Hospital, London, UK), Dr. W.C.G. Overweg-Plandsoen (Academisch Ziekenhuis bij de Universiteit van Amsterdam, Netherlands), Dr. Dorothea Möslinger (Universitätsklinik für Kinder und Jugendliche, Wien, Austria), Dr. Anne-Margret Wingen (Universitätsklinikum Essen, Germany) and Dr. Ina Knerr (University Children’s Hospital, Dublin, Ireland) for sending patient material and data. We thank the Diamond Light Source for access to the beamlines and Carmen Diez-Fernandez for critical comments on the manuscript.

## Materials and Methods

### Reagents

Unless otherwise noted, all compounds were obtained from Sigma-Aldrich (Buchs SG, Switzerland) and were reagent grade or better.

### Purification, crystallization and structure determination of hMCEE

hMCEE was cloned, expressed and purified as previously described (15). Briefly, wt and variant proteins were expressed in *Escherichia coli* BL21(DE3)R3-Rosetta cells from 3-6 L of Terrific Broth culture. Cell pellets were lysed by high pressure homogenizer and centrifuged at 35,000 x*g*. The clarified cell extract was incubated with 2.5 ml of Ni-NTA resin pre-equilibrated with lysis buffer (50 mM HEPES pH 7.5, 500 mM NaCl, 10 mM Imidazole, 5% Glycerol, 0.5 mM TCEP). The column was washed with 100 ml Binding Buffer (50 mM HEPES pH 7.5, 500 mM NaCl, 5% glycerol, 10 mM Imidazole, 0.5 mM TCEP), 50 ml Wash Buffer (50 mM HEPES pH 7.5, 500 mM NaCl, 5% glycerol, 40 mM Imidazole, 0.5 mM TCEP) and eluted with 15 ml of Elution Buffer (50 mM HEPES pH 7.5, 500 mM NaCl, 5% glycerol, 250 mM Imidazole, 0.5 mM TCEP). The eluant fractions were concentrated to 5 ml and applied to a Superdex 200 16/60 column pre-equilibrated in GF Buffer (10 mM HEPES pH 7.5, 500 mM NaCl, 0.5 mM TCEP, 5% glycerol). Eluted protein fractions were incubated with 1:20 mol:mol TEV protease overnight at 4°C. The next day sample was passed through 0.5 ml Ni-NTA pre-equilibrated with GF Buffer and washed 1 ml of GF Buffer. Flow-through and wash were pooled and concentrated to 10-15 mg/ml.

Crystals of hMCEE_WT_ were grown by vapour diffusion in sitting drop at 20°C. A sitting drop consisting of 75 nl protein and 75 nl well solution was equilibrated against well solution containing 30% (v/v) low molecular weight PEG smear (16) and 0.1 M Tris pH 8.5. Crystals were mounted in the presence of 25% (v/v) ethylene glycol and flash-cooled in liquid nitrogen. Crystals of hMCEE_R143C_ were grown by vapour diffusion in sitting drop at 20°C. A sitting drop consisting of 50 nl protein and 100 nl well solution was equilibrated against well solution containing 20% PEG4000, 10% 2-propanol and 0.1M HEPES pH 7.5. Crystals were mounted in the presence of 25% (v/v) ethylene glycol and flash-cooled in liquid nitrogen. Crystallization and diffraction data are given in Table 2. The structure of hMCEE_WT_ was solved by molecular replacement using PHASER (17), with *P. shermanii* MCEE structure (PDB 1JC5) as search model. The structure of hMCEE_R143C_ was solved by molecular replacement with hMCEE wt as search model. Iterative cycles of refinement and manual model building using COOT (18), REFMAC5 (19) and phenix.refine (20) (PMID: 22505256).

### Size Exclusion Chromatography

Size exclusion chromatography was performed as described in (21).

### Nano-Differential Scanning Fluorimetry

Melting curves for the wt protein and mutants were obtained via detection of changes in light scattering using the Prometheus NT.48. Protein concentration was kept at 100 µM in 50 mM HEPES pH 7.5, 500 mM NaCl, 0.5 mM TCEP, 5% glycerol and a melt gradient of 1 degree per minute 20°C to 95°C was used.

### Patient Characterization (genotyping, propionate incorporation)

Skin fibroblasts were taken from patients with biochemical and clinical evidence of methylmalonic aciduria, referred to our institution for diagnostic purposes. The study has been approved by the ethics commission of the Canton of Zurich, Switzerland (application no. 2014-0211). Genomic DNA and RNA extraction, as well as sequencing and propionate incorporation were performed as previously described (21). The nomenclature of the mutations is based on the cDNA sequence NM_032601.3. Nucleotide numbering uses +1 as the A of the ATG translation initiation codon in the reference sequence, with the initiation codon as codon 1.

### Transfection, Immunoblotting and Enzymatic Assay

A DNA fragment encompassing the entire coding sequence of wild-type MCEE was cloned into pcDNA3-CT10HF-LIC using LIC cloning. The sequence was as given by NM_032601.3, except for c.311T>G (p.Lue104Arg), whereby c.311G is the more common allele, (12). Site-directed mutagenesis was carried out on this construct using the QuikChange site-directed mutagenesis kit (Stratagene, La Jolla, CA) as described in the manufacturer’s instructions, using forward and reverse primers (Microsynth, Balgach, Switzerland) and confirmed by Sanger sequencing. Control (BJ, CRL-2522, ATCC) or patient (carrying homozygous MCEE mutation c.139C>T, p.Arg47*) fibroblasts were transiently transfected with 10 µg wild-type or mutant MCEE constructs, with or without 10 µg MUT in pTracer (22) for the enzymatic assay, using electroporation (23). Cells were grown in Dulbecco’s Modified Eagle Medium (Gibco) supplemented with 10% fetal bovine serum (Gibco) and antibiotics (GE Healthcare), as previously described (24) and harvested by trypsinization 48 hr after electroporation, washed twice with HBSS (Gibco) and either frozen at −20°C or processed directly.

Western Blot analysis was performed on fresh or frozen cell pellets essentially as described in (22), with the exception that monoclonal anti-flagM2 (1:4,000; Sigma–Aldrich) and anti-β-Actin (at 1:5,000; Sigma–Aldrich) were used.

For the enzymatic assay, fresh or frozen cell pellet was lysed in buffer including 12 mM Tris-HCl, pH 8.0 and 1mM DTT with sonication twice at amplitude 1.5 for 15 seconds using the microprobe of an XL-2000 sonicator (Microson, Qsonica Newtown, CT). Following lysis, protein concentration was determined by the Lowry method. The incubation mixture contained 300-500 µg cell protein, reaction buffer (100 mM Tris-HCl, pH 8.0; 6 mM MgCl_2_; 3.15 mM ATP; 100 mM KCl; 3 mM DTT), 1 mM propionyl-CoA mix (1 mM propionyl-CoA and 7 µM [^14^C]-propionyl-CoA at 55 mCi/mmol, Anawa, Switzerland) and 50 uM adenosylcobalamin and reaction time was 60 minutes at 37°C. The reaction was stopped by adding 0.5M KOH, samples were then re-incubated for 15 minutes at 37°C to hydrolyze CoA derivatives, neutralized by adding 0.5 N perchloric acid (Merck), and spiked with 0.05% succinic acid, 0.018% methylmalonic acid and 0.018% propionic acid in order to visualize peaks during HPLC separation. Samples were centrifuged to remove precipitate, and supernatant was injected into an Aminex HPX-87H Ion Exclusion column (300 × 7.8 mm2; H-form, 9 μm, Bio-Rad), and organic acids separated by elution with 0.5 mM H_2_SO_4_ at 30°C using a flow rate of 0.4 ml/min. Retention times were 13-15 minutes for methylmalonic acid, 17-19 minutes for succinic acid, and 26-28 minutes for propionic acid (supplemental Fig. S5 and 6), visualized at 210 nm using a UV detector. Fractions covering the methylmalonic acid and succinic acid peaks were collected and [^14^C]-methylmalonic acid and [^14^C]-succinic acid quantified in a Tri-Carb C1 900TR scintillation spectrometer (PerkinElmer,Waltham, MA, USA) with OptiphaseHiSafe 2 counting cocktail (PerkinElmer).

## References

[1] Dobson, C. M., Gradinger, A., Longo, N., Wu, X., Leclerc, D., Lerner-Ellis, J., Lemieux, M., Belair, C., Watkins, D., Rosenblatt, D. S., and Gravel, R. A. (2006) Homozygous nonsense mutation in the MCEE gene and siRNA suppression of methylmalonyl-CoA epimerase expression: a novel cause of mild methylmalonic aciduria. Mol Genet Metab 88, 327–333

[2] Bikker, H., Bakker, H. D., Abeling, N. G., Poll-The, B. T., Kleijer, W. J., Rosenblatt, D. S., Waterham, H. R., Wanders, R. J., and Duran, M. (2006) A homozygous nonsense mutation in the methylmalonyl-CoA epimerase gene (MCEE) results in mild methylmalonic aciduria. Hum Mutat 27, 640–643

[3] Gradinger, A. B., Belair, C., Worgan, L. C., Li, C. D., Lavallee, J., Roquis, D., Watkins, D., and Rosenblatt, D. S. (2007) Atypical methylmalonic aciduria: frequency of mutations in the methylmalonyl CoA epimerase gene (MCEE). Hum Mutat 28, 1045

[4] Mazzuca, M., Maubert, M. A., Damaj, L., Clot, F., Cadoudal, M., Dubourg, C., Odent, S., Benoit, J. F., Bahi-Buisson, N., Christa, L., and de Lonlay, p. (2015) Combined Sepiapterin Reductase and Methylmalonyl-CoA Epimerase Deficiency in a Second Patient: Cerebrospinal Fluid Polyunsaturated Fatty Acid Level and Follow-Up Under L-DOPA, 5-HTP and BH4 Trials. JIMD Rep 22, 47–55

[5] Waters, p. J., Thuriot, F., Clarke, J. T., Gravel, S., Watkins, D., Rosenblatt, D. S., and Levesque, S. (2016) Methylmalonyl-coA epimerase deficiency: A new case, with an acute metabolic presentation and an intronic splicing mutation in the MCEE gene. Mol Genet Metab Rep 9, 19–24

[6] Abily-Donval, L., Torre, S., Samson, A., Sudrie-Arnaud, B., Acquaviva, C., Guerrot, A. M., Benoist, J. F., Marret, S., Bekri, S., and Tebani, A. (2017) Methylmalonyl-CoA Epimerase Deficiency Mimicking Propionic Aciduria. Int J Mol Sci 18

[7] Armstrong, R. N. (2000) Mechanistic diversity in a metalloenzyme superfamily. Biochemistry 39, 13625–13632

[8] He, p., and Moran, G. R. (2011) Structural and mechanistic comparisons of the metal-binding members of the vicinal oxygen chelate (VOC) superfamily. J Inorg Biochem 105, 1259–1272

[9] Meng, E. C., and Babbitt, p. C. (2011) Topological variation in the evolution of new reactions in functionally diverse enzyme superfamilies. Curr Opin Struct Biol 21, 391–397

[10] McCarthy, A. A., Baker, H. M., Shewry, S. C., Patchett, M. L., and Baker, E. N. (2001) Crystal structure of methylmalonyl-coenzyme A epimerase from P. shermanii: a novel enzymatic function on an ancient metal binding scaffold. Structure 9, 637–646

[11] Shi, L., Gao, p., Yan, X. X., and Liang, D. C. (2009) Crystal structure of a putative methylmalonyl-coenzyme A epimerase from Thermoanaerobacter tengcongensis at 2.0 A resolution. Proteins 77, 994–999

[12] Lek, M., Karczewski, K. J., Minikel, E. V., Samocha, K. E., Banks, E., Fennell, T., O’Donnell-Luria, A. H., Ware, J. S., Hill, A. J., Cummings, B. B., Tukiainen, T., Birnbaum, D. P., Kosmicki, J. A., Duncan, L. E., Estrada, K., Zhao, F., Zou, J., Pierce-Hoffman, E., Berghout, J., Cooper, D. N., Deflaux, N., DePristo, M., Do, R., Flannick, J., Fromer, M., Gauthier, L., Goldstein, J., Gupta, N., Howrigan, D., Kiezun, A., Kurki, M. I., Moonshine, A. L., Natarajan, p., Orozco, L., Peloso, G. M., Poplin, R., Rivas, M. A., Ruano-Rubio, V., Rose, S. A., Ruderfer, D. M., Shakir, K., Stenson, p. D., Stevens, C., Thomas, B. P., Tiao, G., Tusie-Luna, M. T., Weisburd, B., Won, H. H., Yu, D., Altshuler, D. M., Ardissino, D., Boehnke, M., Danesh, J., Donnelly, S., Elosua, R., Florez, J. C., Gabriel, S. B., Getz, G., Glatt, S. J., Hultman, C. M., Kathiresan, S., Laakso, M., McCarroll, S., McCarthy, M. I., McGovern, D., McPherson, R., Neale, B. M., Palotie, A., Purcell, S. M., Saleheen, D., Scharf, J. M., Sklar, p., Sullivan, p. F., Tuomilehto, J., Tsuang, M. T., Watkins, H. C., Wilson, J. G., Daly, M. J., MacArthur, D. G., and Exome Aggregation, C. (2016) Analysis of protein-coding genetic variation in 60,706 humans. Nature 536, 285–291

[13] Schymkowitz, J., Borg, J., Stricher, F., Nys, R., Rousseau, F., and Serrano, L. (2005) The FoldX web server: an online force field. Nucleic Acids Res 33, W382–388

[14] Grunert, S. C., Stucki, M., Morscher, R. J., Suormala, T., Burer, C., Burda, p., Christensen, E., Ficicioglu, C., Herwig, J., Kolker, S., Moslinger, D., Pasquini, E., Santer, R., Schwab, K. O., Wilcken, B., Fowler, B., Yue, W. W., and Baumgartner, M. R. (2012) 3-methylcrotonyl-CoA carboxylase deficiency: clinical, biochemical, enzymatic and molecular studies in 88 individuals. Orphanet J Rare Dis 7, 31

[15] Hamed, R. B., Gomez-Castellanos, J. R., Sean Froese, D., Krysztofinska, E., Yue, W. W., and Schofield, C. J. (2016) Use of Methylmalonyl-CoA Epimerase in Enhancing Crotonase Stereoselectivity. Chembiochem 17, 471–473

[16] Chaikuad, A., Knapp, S., and von Delft, F. (2015) Defined PEG smears as an alternative approach to enhance the search for crystallization conditions and crystal-quality improvement in reduced screens. Acta Crystallogr D Biol Crystallogr 71, 1627–1639

[17] McCoy, A. J. (2007) Solving structures of protein complexes by molecular replacement with Phaser. Acta Crystallogr D Biol Crystallogr 63, 32–41

[18] Emsley, p., Lohkamp, B., Scott, W. G., and Cowtan, K. (2010) Features and development of Coot. Acta Crystallogr D Biol Crystallogr 66, 486–501

[19] Murshudov, G. N., Vagin, A. A., and Dodson, E. J. (1997) Refinement of macromolecular structures by the maximum-likelihood method. Acta Crystallogr D Biol Crystallogr 53, 240–255

[20] Adams, p. D., Afonine, p. V., Bunkoczi, G., Chen, V. B., Davis, I. W., Echols, N., Headd, J. J., Hung, L. W., Kapral, G. J., Grosse-Kunstleve, R. W., McCoy, A. J., Moriarty, N. W., Oeffner, R., Read, R. J., Richardson, D. C., Richardson, J. S., Terwilliger, T. C., and Zwart, p. H. (2010) PHENIX: a comprehensive Python-based system for macromolecular structure solution. Acta Crystallogr D Biol Crystallogr 66, 213–221

[21] Plessl, T., Burer, C., Lutz, S., Yue, W. W., Baumgartner, M. R., and Froese, D. S. (2017) Protein destabilization and loss of protein-protein interaction are fundamental mechanisms in cblA-type methylmalonic aciduria. Hum Mutat 38, 988–1001

[22] Forny, p., Froese, D. S., Suormala, T., Yue, W. W., and Baumgartner, M. R. (2014) Functional characterization and categorization of missense mutations that cause methylmalonyl-CoA mutase (MUT) deficiency. Hum Mutat 35, 1449–1458

[23] Baumgartner, M. R., Almashanu, S., Suormala, T., Obie, C., Cole, R. N., Packman, S., Baumgartner, E. R., and Valle, D. (2001) The molecular basis of human 3-methylcrotonyl-CoA carboxylase deficiency. J Clin Invest 107, 495–504

[24] Suormala, T., Baumgartner, M. R., Coelho, D., Zavadakova, p., Kozich, V., Koch, H. G., Berghauser, M., Wraith, J. E., Burlina, A., Sewell, A., Herwig, J., and Fowler, B. (2004) The cblD defect causes either isolated or combined deficiency of methylcobalamin and adenosylcobalamin synthesis. J Biol Chem 279, 42742–42749

