## Supplementary material for "Genetic, structural, and functional analysis of mutations causing methylmalonyl-CoA epimerase deficiency"

^3^Present address: Institute of Pharmaceutical Chemistry & Structural Genomics Consortium, BMLS, Goethe-University Frankfurt, 60438 Frankfurt, Germany

^4^Present address: Astex Pharmaceuticals, 436 Cambridge Science Park, Milton Road, Cambridge, CB4 0QA, UK

^‡^ equal contribution

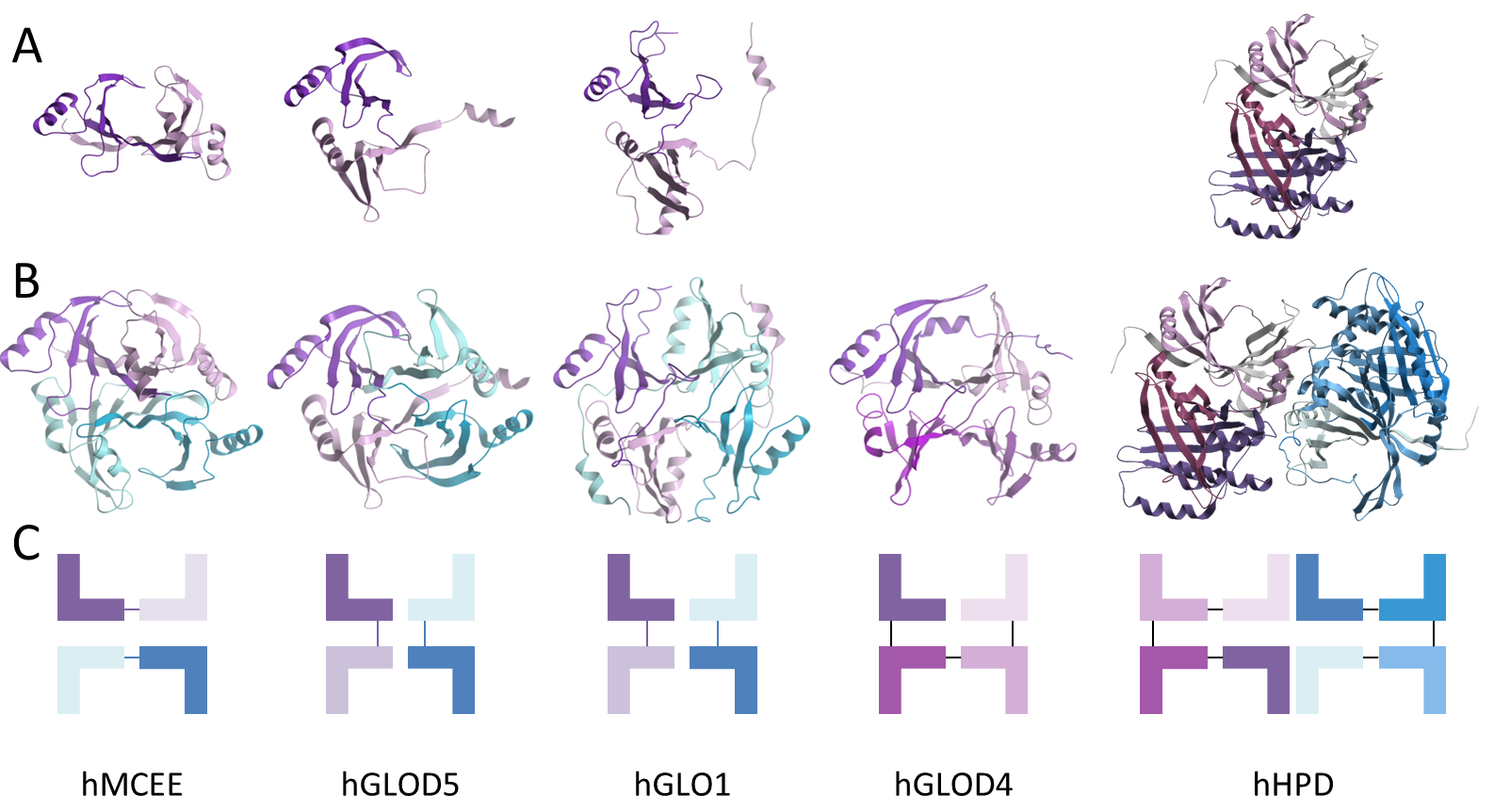

**Supplementary Figure 1.** **Modular arrangement of human VOC family proteins.** **A.** Monomeric subunits. **B.** Physiological arrangement. **C.** Schematic arrangement of each protein. For the bottom row, each “L” represents one GLOD domain. PDB IDs: hMCEE (3RMU), hGLOD5 (3ZW5), hGLO1 (3VW9), hGLOD4 (3ZI1) and hHPD (3ISQ).

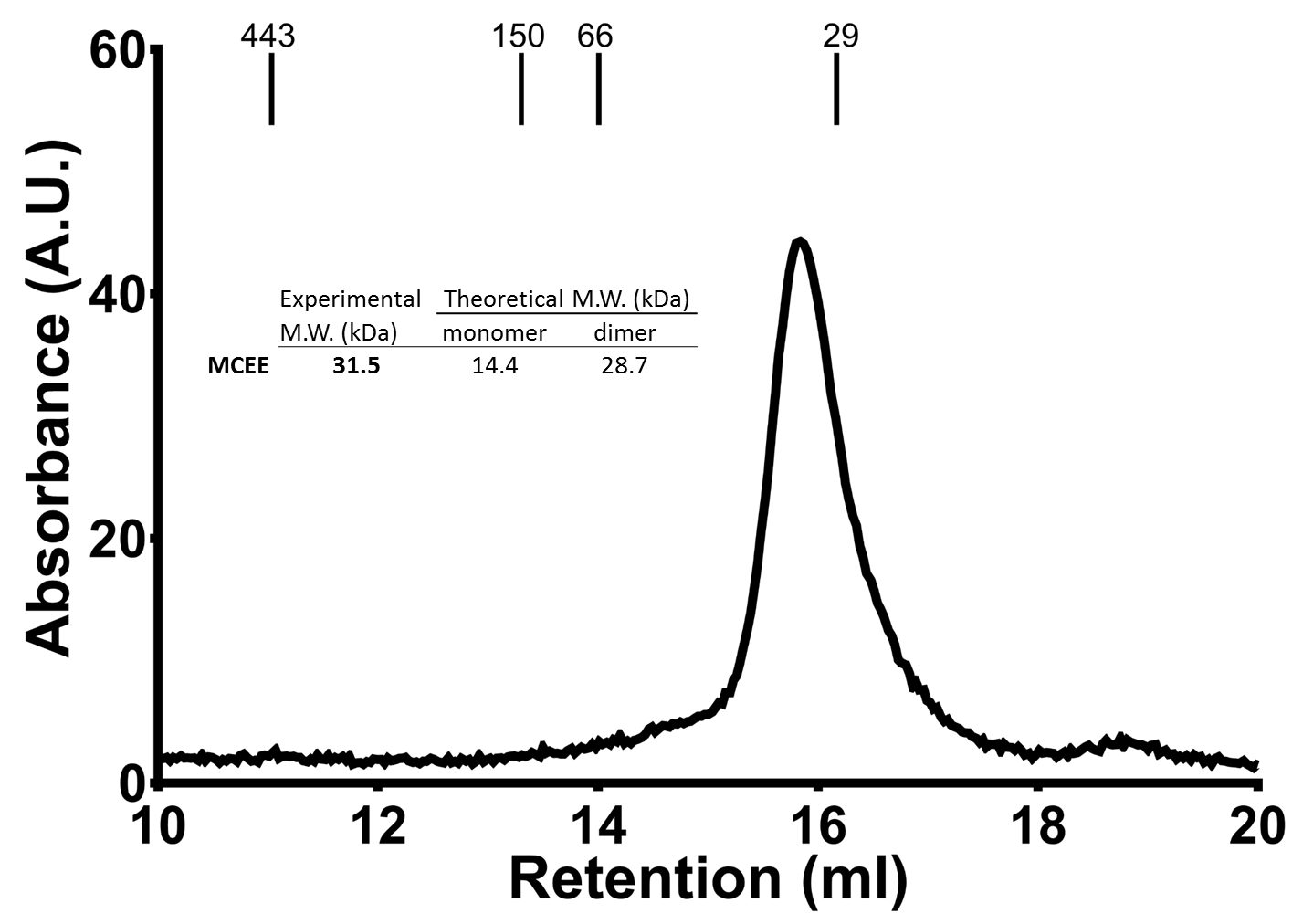

**Supplementary Figure 2. Size-exclusion chromatography of hMCEE.** Absorbance was monitored at 280nm and is given in absorbance units (A.U.). Analytical gel filtration was performed on a Superdex 200 HiLoad 10/30 column (GE Healthcare) pre-equilibrated with 25 mM HEPES pH 7.5, 150 mM NaCl and 5% glycerol. The column was calibrated using carbonic anhydrase (29 kDa), bovine serum albumin (66 kDa), alcohol dehydrogenase (150 kDa), and apoferritin (443 kDa) as protein standards. Experiments were performed with 50 μM of protein. Molecular weight (M.W. in kiloDaltons, kDa) was calculated as outlined in (1).

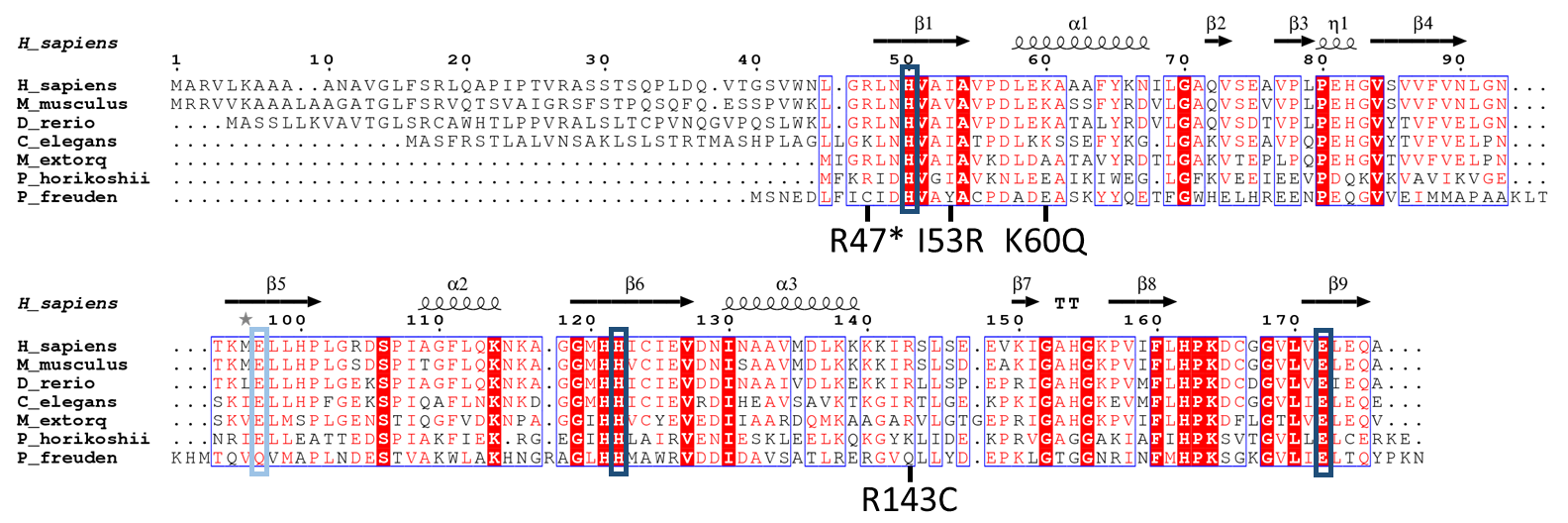

**Supplementary Figure 3. Sequence alignment of MCEE orthologs ranging from human to bacteria.** Sequences used were: H_sapiens, NP_115990.3; M_musculus, NP_082902.1; D_rerio, NP_001018602.1; C_elegans, NP_492120.1; M_extorquens, OHV16813.1; P_horikoshii, WP_048053074.1; P_freuden, WP_055344597.1. Absolutely conserved residues are highlighted in red shading, well-conserved residues in red font. Secondary structure elements from *h*MCEE are indicated at the top of the alignment, *α*-helices as coils, and *β*-sheets as black arrows. Cobalt binding residues are marked in black boxes, the active-site residue Glu97 (hMCEE numbering) in a blue box, and patient mutations are marked below. Alignment was created by multalin (2) and visualized by ESPript (3).

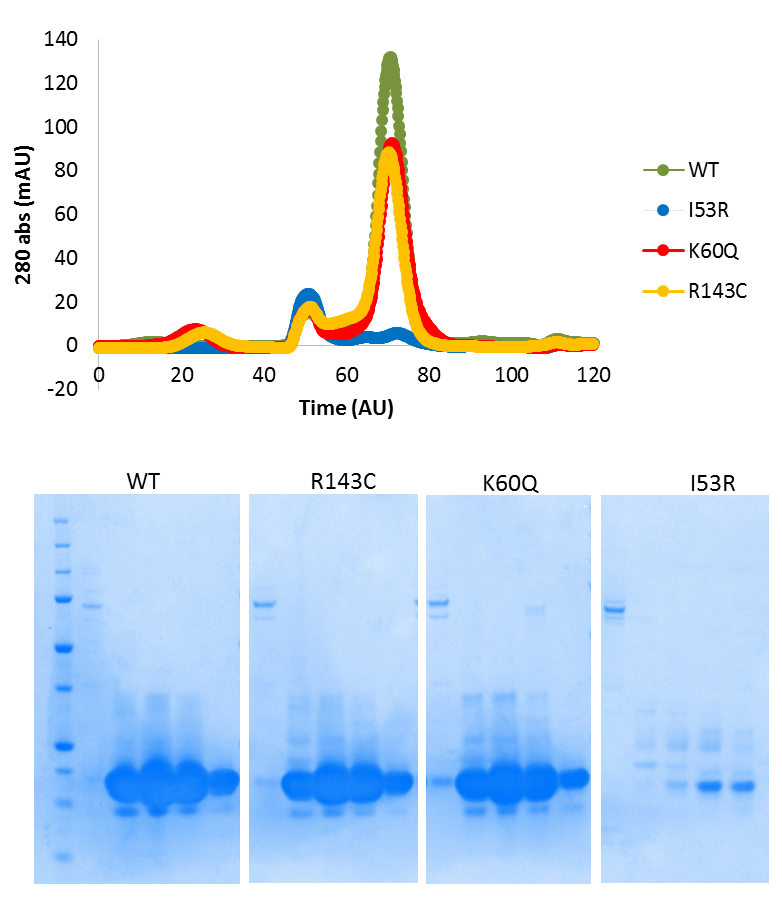

**Supplementary Figure 4.** Gel filtration experiments of WT and mutant enzymes were performed using a Superdex 75 HiLoad 16/60 column pre-equilibrated with 50 mM HEPES pH 7.5, 500 mM NaCl, 0.5 mM TCEP, and 5% glycerol. Lane one of each gel corresponds to the void peak at time point 50 (AU). The remaining 4 lanes correspond to fractions under the main peak between time points 60 and 80 (AU).

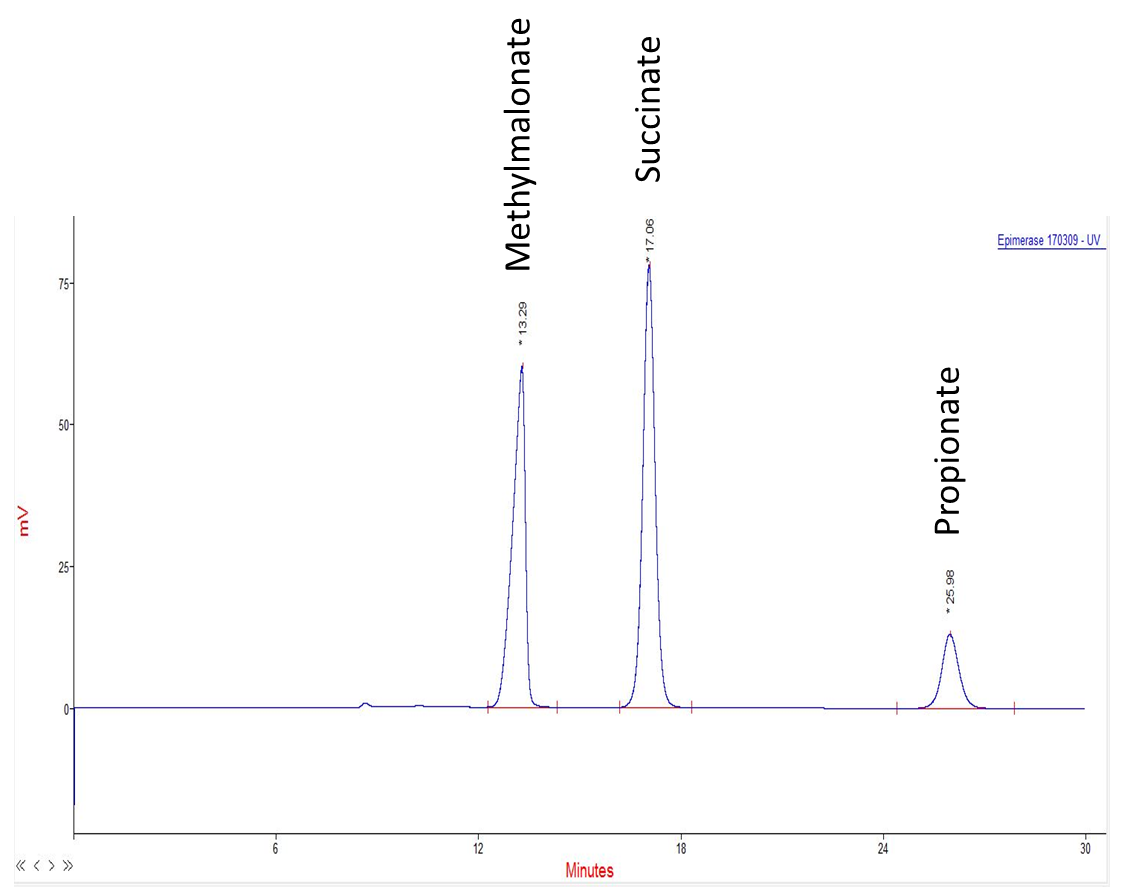

**Supplementary Figure 5. Example of the separation of standards containing methylmalonic acid, succinate and propionic acid by HPLC.**

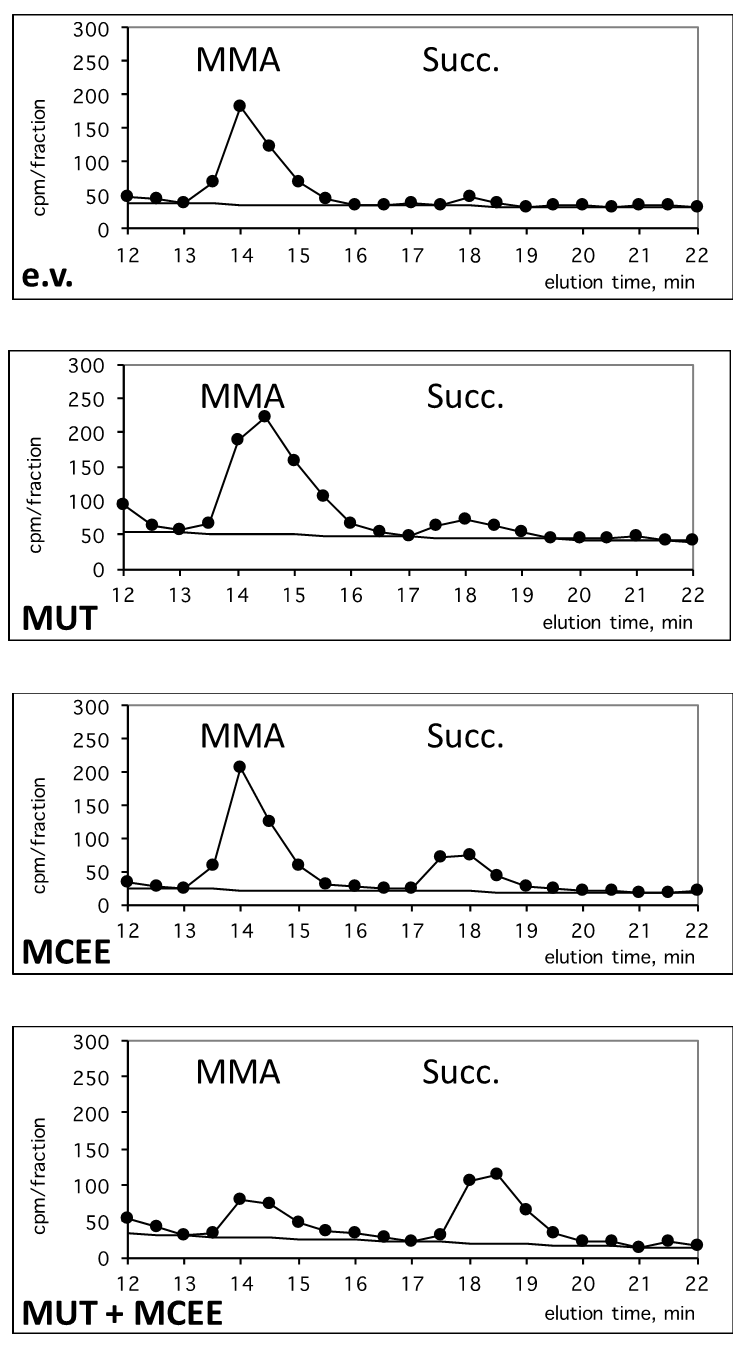

**Supplementary Figure 6. Example of detection of radio-labeled methylmalonic acid (MMA) and succinate (Succ.) following separation and fractionation in an HPLC.** In the bottom left-hand corner of each panel is given which proteins are over-expressed. e.v. is empty vector, MUT is methylmalonyl-CoA mutase. MMA and Succ. are written above the peaks to which they correspond.

**References**

1.	Irvine, G. B. (2001) Determination of molecular size by size-exclusion chromatography (gel filtration). *Curr Protoc Cell Biol* **Chapter 5**, Unit 5 5

2.	Corpet, F. (1988) Multiple sequence alignment with hierarchical clustering. *Nucleic Acids Res* **16**, 10881-10890

3.	Gouet, P., Robert, X., and Courcelle, E. (2003) ESPript/ENDscript: Extracting and rendering sequence and 3D information from atomic structures of proteins. *Nucleic Acids Res* **31**, 3320-3323
